# Genomic analysis reveals the interplay between ABA-GA in determining dormancy duration in groundnut

**DOI:** 10.1101/2025.06.20.660118

**Authors:** D. Khaja Mohinuddin, Sunil S. Gangurde, Hasan Khan, Deekshitha Bomireddy, Vinay Sharma, Priya Shah, U. Nikhil Sagar, Namita Dube, Ramachandran Senthil, B.V. Tembhurne, V. Hanumanth Nayak, A. Amaregouda, Kisan Babu, Kuldeep Singh, Pasupuleti Janila, Baozhu Guo, Boshou Liao, Rajeev K. Varshney, Manish K. Pandey

## Abstract

Groundnut is an important leguminous crop however; its productivity and seed quality are frequently reduced due to lack of fresh seed dormancy (FSD). To address this challenge, a mini-core collection of 184 accessions was phenotyped to identify donors in each agronomic type, in addition to analysing data on whole genome re-sequencing and multi-season phenotypic evaluations to identify stable marker-trait associations (MTAs) associated with FSD. Phenotypic analysis revealed substantial variability in dormancy durations, with days to 50% germination (DFG) ranging from 1 to 30 days. Multi-locus genome-wide association studies (ML-GWAS) identified 27 MTAs in individual seasons and 12 MTAs in pooled seasons data, respectively. Key candidate genes identified included *Cytochrome P450 superfamily proteins, protein kinase superfamily proteins,* and *MYB transcription factors* involved in the Abscisic acid (ABA) pathway, as well as *F-box interaction domain proteins, ATP-binding ABC transporters,* associated with the Gibberellic acid (GA) pathway. SNP-based KASP (Kompetitive Allele-Specific Polymerase chain reaction) markers for 12 SNPs were developed and validated, of these 6 markers (snpAH00577, snpAH00580, snpAH00582, snpAH00585, snpAH00586 and snpAH00588) showed polymorphism between dormant and non-dormant lines. Incorporating favourable dormant alleles into breeding strategies could enable the development of high-yielding cultivars with a dormancy period of 2-3 weeks.

## 1. Introduction

Groundnut (*Arachis hypogaea* L.) is a globally important legume crop with significant economic and nutritional value. It is well-known for its rich nutritional content, offering a valuable source of protein, healthy unsaturated fatty acids, and essential vitamins [1]. In 2023, the Food and Agriculture Organization (FAO) reported that the worldwide groundnut harvest area surpassed 32.7 million hectares, leading to a total production of over 54 million tons. Seeds are absolutely vital for plant growth and development, acting as the primary carriers of genetic traits, passing on essential characteristics from one generation to the next [2]. This genetic transfer is vital for producing high-quality crops in the field.

Cultivated groundnut can be classified into four agronomic types-Spanish bunch, Valencia bunch, Virginia bunch and Virginia runner, they belong to different subspecies (spp. *hypogaea* and *fastigiata*) - based on developmental patterns, flower arrangement, kernel and, pod traits [3]. Valencia bunch and Spanish bunch genotypes had shorter maturity duration and no seed dormancy (SD), while the Virginia bunch and Virginia runner genotypes often had longer maturity duration and have seeds with variable dormancy durations [4]. Among these, Spanish groundnut varieties are mainly grown in Africa and semi-arid regions of Asia, and they make up 60% of the world’s groundnut production. However, early rainfall before harvest can lead to pre-harvest sprouting (PHS), where seeds germinate prematurely. This can cause a 10–20% reduction in yield, with Spanish cultivars being particularly vulnerable, sometimes experiencing losses of up to 50%. These losses not only reduce yield but also increase the risk of pathogen infections, elevate the chances of aflatoxin contamination, lower market prices, and degrade kernel quality [5].

SD is characterized by the inability of a viable seed to germinate even under favourable conditions [6]. The concept of “germination” refers to the metabolic activation of a seed upon absorbing water, which causes the radicle to emerge through the seed coat, this process is also known as “chitting.” According to seed technology, germination is the process by which a seedling emerges and grows from an embryo, exhibiting vital components that suggest the seedling has the capacity to become a regular seedling in the right circumstances [7]. SD is regulated by a complex interplay of hormonal, molecular, and environmental factors. Across many plant species, hormonal control of dormancy follows a conserved pattern, with ABA functioning as the primary regulator. ABA is essential for both the initiation and maintenance of dormancy. In contrast, GA facilitate dormancy release and promote germination [8]. In modern groundnut cultivation, achieving an optimal dormancy period of 2–3 weeks requires the appropriate combination of genetic factors. Groundnut breeding programs increasingly rely on genomics-assisted breeding (GAB) strategies, utilizing diagnostic markers and advanced molecular tools to enhance key traits such as oleic acid content and disease resistance [9,10].

To mitigate the adverse effects of PHS, the introduction of FSD lasting 2-3 weeks has emerged as a viable solution. This allows farmers to postpone harvesting during unexpected rainfall, hence minimizing losses from PHS. Therefore, a major objective for plant breeders is to develop high-yielding cultivars with the dormancy trait, since it improves sustainability and resilience in groundnut cultivation [5]. Various studies have demonstrated that the SD is influenced by many factors, including internal (embryo, seed coat, and endogenous inhibitors) [11] and environmental factors (temperature, light and, air etc.) [12].

To uncover genetic variants that influence complex behaviours, GWAS have been utilized to explore potential links between specific genotypes and observable traits. These studies help to bridge the gap between genetic information and its practical applications by identifying the genetic factors that influence various traits and behaviors [13]. This approach has gained recognition as a powerful method for identifying QTLs and genes associated with complex traits, leveraging historical recombination crossovers within large natural populations [14,15]. By tapping into the genetic diversity present in these populations, we can uncover valuable insights into the genetic underpinnings of traits that are often influenced by multiple factors. Unlike QTL mapping, which is constrained by the genetic diversity of the parental lines used in specific crosses, GWAS leverages natural populations that exhibit a wide range of genetic variation, enabling the identification of a broader spectrum of alleles linked to complex traits [16], although are less useful to detecting very rare alleles for a trait. This approach enhances mapping accuracy by utilizing historical recombinations and linkage disequilibrium across diverse germplasm, facilitating the discovery of multiple quantitative trait nucleotides (QTNs) that contribute to the desired traits [17]. Additionally, GWAS is adept at identifying small-effect QTLs that might be missed in biparental populations, offering a more comprehensive view of the genetic architecture underlying various traits.

GWAS have shown to be a useful method for identifying MTAs related to SD and PHS resistance in crops like wheat and rice. For instance, in wheat significant MTAs and putative candidate genes have identified such as ABA-responsive proteins, protein kinases, and MAP-kinase-like proteins [18]. Similarly, studies in rice have found important QTLs linked to PHS resistance and SD, proving the efficacy of GWAS across different germplasm lines [19]. For instance, in rice, the *GA20-oxidase* gene has been identified within the QTL region that controls PHS [20]. Similarly, in barley, genes such as *mitogen-activated protein kinase kinase 3* (MKK3) and *alanine aminotransferase (AlaAT)* have been linked to SD regulation [21]. In wheat, the mother of *FT* and *TFL1 (MFT)* and *Phs1* genes have also been recognized as key regulators of SD [21,22]. While PHS poses a significant challenge in groundnut cultivation, research efforts to map the relevant FSD QTLs have been limited. These findings highlighted potential of GWAS to expand the understanding of genetic factors governing seed dormancy and create the way for targeted breeding methods intended to enhance crop resilience and quality [23,24]. This highlights a gap in understanding the genetic basis of PHS resistance in groundnuts compared to other crops.

Several statistical models have been employed for association mapping, utilizing various methodologies [25,26]. Traditional single-locus genome scans using ordinary mixed models often fall short in accounting for loci with large effects. As a solution, multi-locus genome-wide association study (ML-GWAS) models have been proposed [27,28]. These ML-GWAS models are recognized for their efficiency and reliability in mapping genomic regions, as they estimate the effects of all markers simultaneously. Unlike single-locus genome-wide association studies (SL-GWAS), ML-GWAS do not necessitate stringent multiple testing corrections, which often lead to the dismissal of significant associations [29]. Furthermore, ML-GWAS models exhibit greater power in detecting significant marker-trait associations compared to single-locus approaches [26].

To address the existing knowledge gaps and leverage advancements in association mapping analysis. This study employed GWAS models on a diverse mini-core collection of groundnut to identify significant MTAs related to FSD. By integrating whole genome re-sequencing data with multi-season phenotypic data on a mini-core collection, we aimed to uncover candidate genes that regulate FSD, ultimately facilitating the development and validation of molecular markers for use in breeding. This approach enhances our understanding of the genetic basis of FSD and contributes to the development of resilient groundnut varieties.

## 2. MATERIAL AND METHODS

### 2.1 Plant material and phenotyping for fresh seed dormancy

In present study, the experimental material comprised of 184 accessions from the groundnut mini-core collection developed at ICRISAT [30]. This mini-core set represents 1.29% of the full collection and 10.8% of the core collection, respectively. The mini-core subset attempts to represent the overall genetic diversity of the ICRISAT gene bank collection. The 184 accessions in the mini-core set represents different botanical groups of groundnut, including 33 accessions of Virginia runner, 48 of Virginia bunch, 57 Spanish bunch and 35 Valencia bunch. The significant genetic diversity of this mini-core collection makes it an informative panel for genetic dissection of complex traits in groundnut. It can also be utilized for molecular characterization, enabling the selection of parents and maximizing diversity in groundnut breeding programs to enhance the heritability and genetic gain (**Figure 1a**).

**Figure 1.**
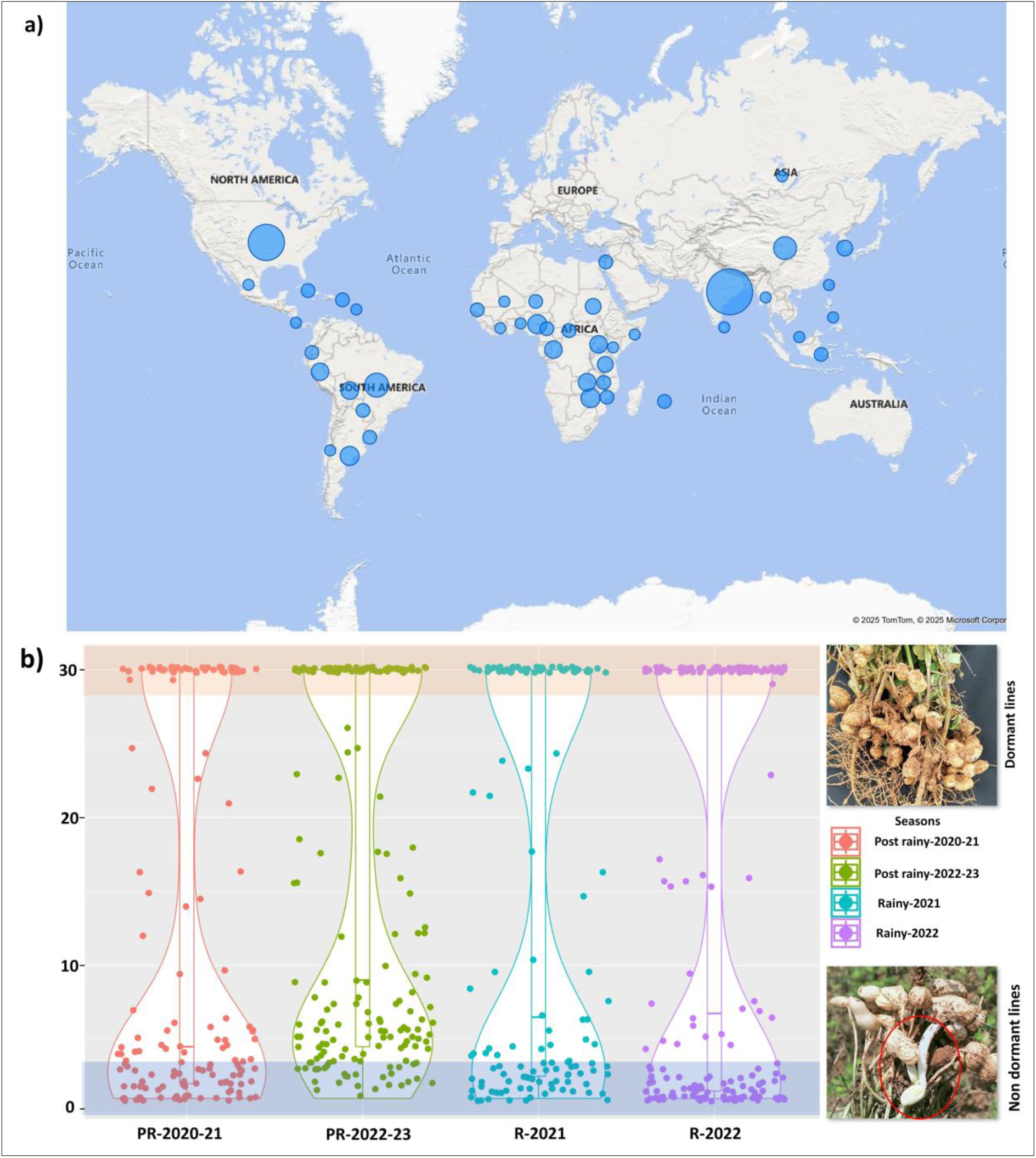
Phenotypic variability for fresh seed dormancy in the mini-core accessions. a) The world map illustrates the geographical distribution of the groundnut mini-core collection globally. Countries are represented by circles of varying sizes, which indicate the presence and diversity of accessions. Larger circles denote a higher number of accessions, while smaller circles represent fewer accessions. **b)** The violin plot displays the dormancy characteristics of groundnut lines over four seasons: Post rainy 2020-21, Post rainy 2022-23, Rainy 2021, and Rainy 2022. Each violin represents the distribution of data points for that season, with higher points indicating dormancy and lower points showing non-dormant lines. The shaded areas within the plot indicate the density of data points.

The groundnut mini-core collection was sown during two post rainy seasons (PR2020-2021 and PR2022-2023) and one rainy season (R-2022) at ICRISAT, Hyderabad, 17◦31′48.00′′ N latitude, 78◦16′12.00′′ E longitude, and an altitude of 545 m. Additionally, during the 2021 rainy season, the accessions were also planted at the SV Agricultural College farm in Tirupati, situated at 13°54’ N latitude, 79°54’ E longitude, and 182.9 m altitude. The experiment was conducted in a randomized block design (RBD), two replications, and with two plant and row spacing of 10cm and 60cm, respectively, under recommended agronomic conditions for groundnut.

For evaluation of FSD, fully matured seeds were freshly harvested and evaluated using an in-vitro germination assay[3]. For each accession, 20 seeds with uniform size were selected from two replicates. Each accession was treated with fungicides (Mancozeb and Carbendazim) and placed on moist germination paper in sterilized petri dishes. Plates were stored in total darkness and watered every 24 hours. Germination data were recorded daily for 30 days, and the number of days required for an accession to reach 50% germination was documented as its dormancy duration.

### 2.2 DNA extraction, sequencing and SNP calling

DNA was extracted from 100 mg of fresh leaf tissue from each accession following the NucleoSpin Plant II kit (Macherey-Nagel, Düren, Germany) [31] instructions. The leaf samples were first homogenized in 500 µL of lysis buffer, and 10 µL of RNase was added to remove RNA contaminants. The sample was placed into a water bath at 65°C for 1 hour. After incubation, the sample was centrifuged for 20 minutes at 6000 rpm, and the supernatant was collected and transferred. The supernatant and 450 µL of binding buffer were placed through a NucleoSpin Plant MN column. The column was centrifuged at 6000 rpm for 1 minute, and the flow-through was discarded. The pellet was washed twice, first with 400 µL of buffer PW1 and then centrifuged. The pellet was then washed with 700 µL buffer PW2 to ensure the pellet was free of any contaminants. A warm elution buffer (65 °C) of 50 µL was added to the membrane filter of the column and incubated for 5 minutes at 65 °C. The elution was performed by centrifugation for 1 minute at 6000 rpm. Following extraction, the concentration and quality of the extracted DNA was assessed using a Nanodrop 8000 Spectrophotometer (Thermo Fisher Scientific, Waltham, MA, USA), and electrophoresis in a 0.8% agarose gel.

Sequencing was performed on Illumina platform and 10X coverage was achieved for each accession. Post sequencing, adapter sequences were trimmed from sequences, and low-quality reads were filtered out. These reads were considered low-quality if they contained more than 20% of bases that were low-quality (quality value ≤ 7) and more than 5% “N” nucleotides; we used SOAP2 to clean the completed sequences to retain only reads with high-quality sequencing data for downstream analysis [32]. Next, we aligned cleaned reads to the reference genome for the cultivated tetraploid variety “Tifrunner” using the same software with parameters of “-m 300 -x 600 -s 35 -l 32 -v 5 -p 4.” After alignment with the reference genome, we calculated the likelihood for all possible genotypes for each sample with SOAPsnp3, which takes into account the maximum likelihood estimates of allele frequencies in the population. To refine the dataset, low-quality variants were filtered based on strict criteria: sequencing depth greater than 10,000 and less than 400, mapping quality scores exceeding 1.5, and overall quality scores below 20. Only loci with estimated allele frequencies that were not equal to 0 or 1 were kept. By the time we were done, we had removed all remaining SNPs with more than 50% missing data per genotype to ensure that all downstream evaluations were derived from high-quality SNP.

### 2.3 Single and multi-locus genome-wide association studies analysis and identification of candidate genes

A total of 5,61,099 SNPs were filtered from WGRS data on 184 mini-core accessions. Minor allele frequency (MAF) was applied at the 0.05 level. A large proportion of heterozygous SNPs were detected, but due to low confidence level in heterozygous state calling, they were considered as missing data, and heterozygosity criteria were applied at a 0.25 level. A working subset of 2,55,144 filtered SNPs with 0.2% heterozygosity, and 0.05% MAF was used for GWAS analysis with bonferroni correction of 0.05% using TASSEL v.5 software [33]. At a stringent level, imputation was not attempted for missing data. To identify MTAs, GWAS analysis was performed using the Genome Association and Prediction Integrated Tool (GAPIT) package using the R software [34].

To pinpoint genetic regions linked to our traits of interest, we conducted a comprehensive GWAS analysis. This involved combining detailed multi-season observations with extensive genetic data – specifically, 255,144 SNPs from our mini-core collection. We used a robust approach, employing both single- and multi-locus statistical models, including CMLM, BLINK, MLMM, and FarmCPU, all within the R statistical environment using the GAPIT package [24]. To ensure reliable results and minimize false positives, we carefully accounted for population structure by incorporating the first three principal components and a kinship matrix in accounting for population structure. The association threshold for significant MTAs was calculated using Bonferroni correction to determine a p-value of 6.7 × 10^-7^ which can be calculated from the negative log conversion of α/n (where n= total SNPs for GWAS analysis) [35]. Candidate genes were identified from 200 kb genomic region, around significant MTAs, with 100 kb upstream, and 100 kb downstream included. Peanut base (https://Peanutbase.org/) was used to identify the candidate genes using GBrowse (cultivated peanut) version 1 using gene ID.

### 2.4 Kompetitive Allele-Specific PCR marker development and validation

Functionally important candidate genes were used for marker development as SNPs were identified from the genomic regions, and the selected SNPs were on 11 different chromosomes adjacent to the target genes. KASP markers were developed and validated using contrasting germplasm lines from the ICRISAT. To ensure the creation of user-friendly and cost-effective markers, 300-bp upstream and 300-bp downstream sequences were incorporated for SNP conversion into KASP markers [36]. Each KASP assay included two allele specific forward primers and one common reverse primer (Intertek Pvt. Ltd.) The validation panel consisted of 37 genotypes, including 16 dormant, 5 moderately dormant, and 16 non-dormant lines (**Supplementary Table 7**). A complete list of the KASP primers used in this study is provided in **Table 2**.

## 3. Results

### 3.1 Phenotypic variation among different agronomic types for fresh seed dormancy in mini-core collection

In this study, FSD was estimated based on the number of days required for an accession to reach 50% germination. The GWAS panel exhibited considerable variation in dormancy duration, ranging from 1 to 30 days. Among the different agronomic types, Virginia bunch and Virginia runner (var. *hypogaea*) accessions displayed the longest dormancy period, lasting 16 to 30 days, whereas Spanish bunch (var. *vulgaris*) and Valencia (var. *fastigiata*) accessions showed dormancy durations ranging from 1 to 25 days. The mini-core collection represented diverse geographic origins, as illustrated in **Figure 1a**. The mean performance and phenotypic distribution of the accessions across multiple seasons were analyzed from two replications, with results depicted in **Figure 1b and Supplementary Table 1**. These findings highlight the significant variability in FSD among the mini-core accessions, providing a strong basis for further genetic studies and association mapping.

Significant variation in SD was observed across seasons among a diverse panel of 184 groundnut mini-core collection. Analysis of variance (ANOVA) indicated highly significant differences among accessions (P < 0.01), highlighting substantial phenotypic diversity within the panel. Violin plots depicted the distribution of trait values for each season, with wider sections of the violins indicated more frequent values, and reflecting the diversity among the accessions (**Figure 1b**). Dormancy periods ranged from 1 to 30 days, with averages durations of 12.54, 14.57, 14.96, and 15.72 days for four seasons, and standard errors of 0.52, 0.41, 3.01, and 2.98, respectively. High heritability (∼90%) further highlighted the genetic control of this trait across the seasons, emphasizing the broad variability within the mini-core collection. This genetic variability laid the foundation for conducting GWAS to identify significant MTAs associated with FSD.

### 3.2 GWAS identified significant MTAs associated with fresh seed dormancy

The analysis of SNP (Single Nucleotide Polymorphism) density across chromosomes revealed substantial variation in SNP distribution. Chromosomes Ah03, Ah19, and Ah04 exhibited the highest SNP densities, with 19001, 15758, and 14193 SNPs, respectively. In contrast, chromosomes Ah08 and Ah07 displayed relatively lower SNP densities. Notably, red-coloured regions indicated areas of high SNP density, particularly on chromosome Ah17. Additionally, white gaps observed on chromosomes Ah14, Ah15, and Ah16 suggest regions with low or undetected SNPs (**Supplementary Figure 1**).

A total 27 MTAs (-log_10_P 6.7) significantly associated with FSD were identified in GWAS analysis (**Table 1**). Among these, 5 MTAs (Ah04_98048339, Ah05_14439200, Ah05_25536212, Ah05_26459808, Ah20_126576288) were consistently identified across at least by two models (**Figure 2a,b; Supplementary Figure 2**). Notably, MTAs on chromosomes Ah05 (Ah05_14439200) and Ah20 (Ah20_126576288) were consistently identified in all four models during PR2020-21 and R2022, with higher phenotypic variance explained (PVE %) of 69.61 and 66.54 respectively. Additionally, 3 MTAs (Ah04_79354879, Ah05_25536212, Ah05_26459808) were identified in at least three models, with the PVE (%) of 37.81, 30.59 and 13.24 during PR2020-21, R2021 and PR2022-23, seasons respectively. Furthermore, 17 MTAs were identified on chromosomes Ah01, Ah03, Ah04, Ah05, Ah11, Ah13, Ah14, Ah18, Ah19, and Ah20, with each chromosome revealing a single model (**Table 1**).

**Table. 1:**
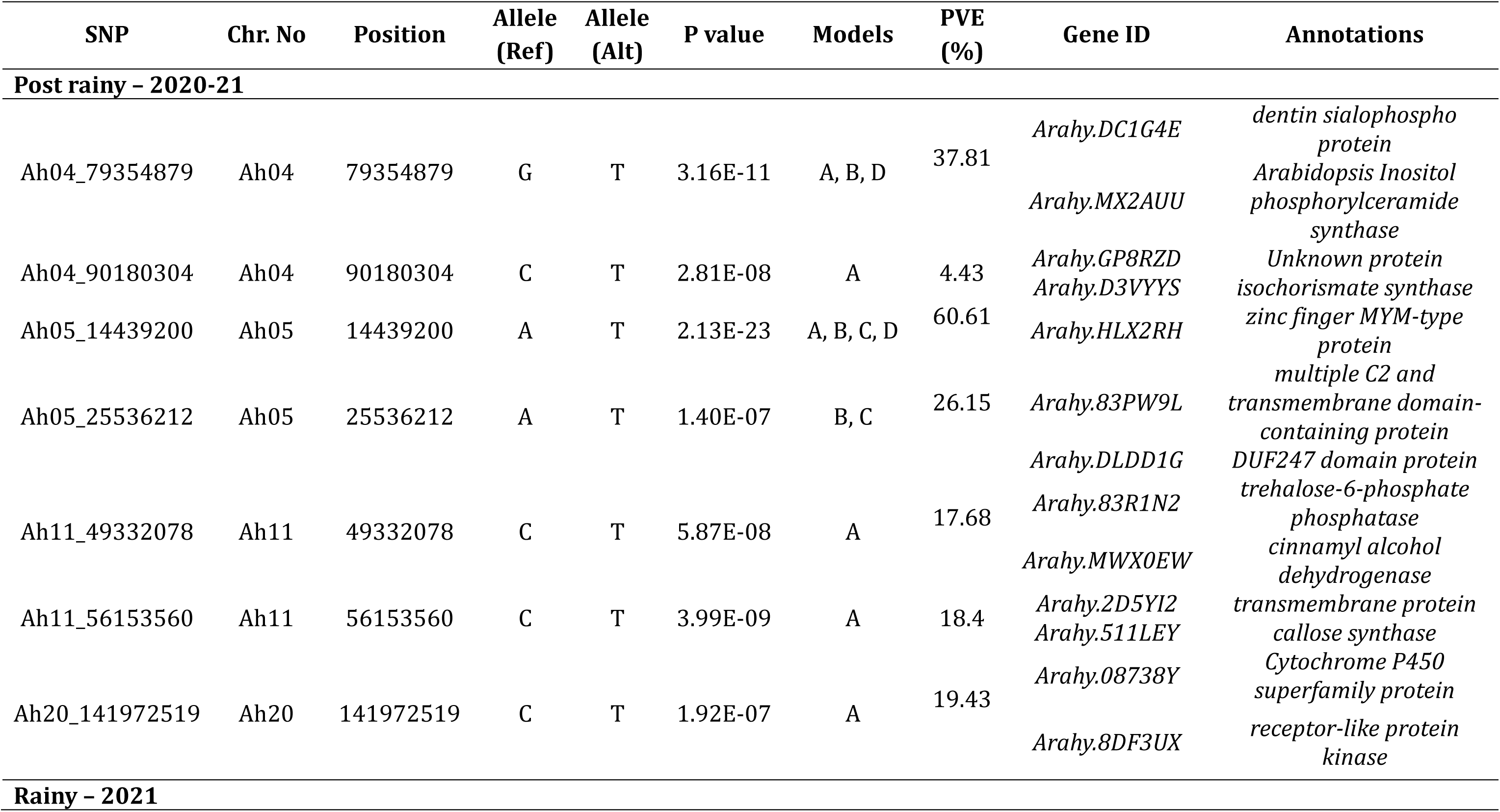
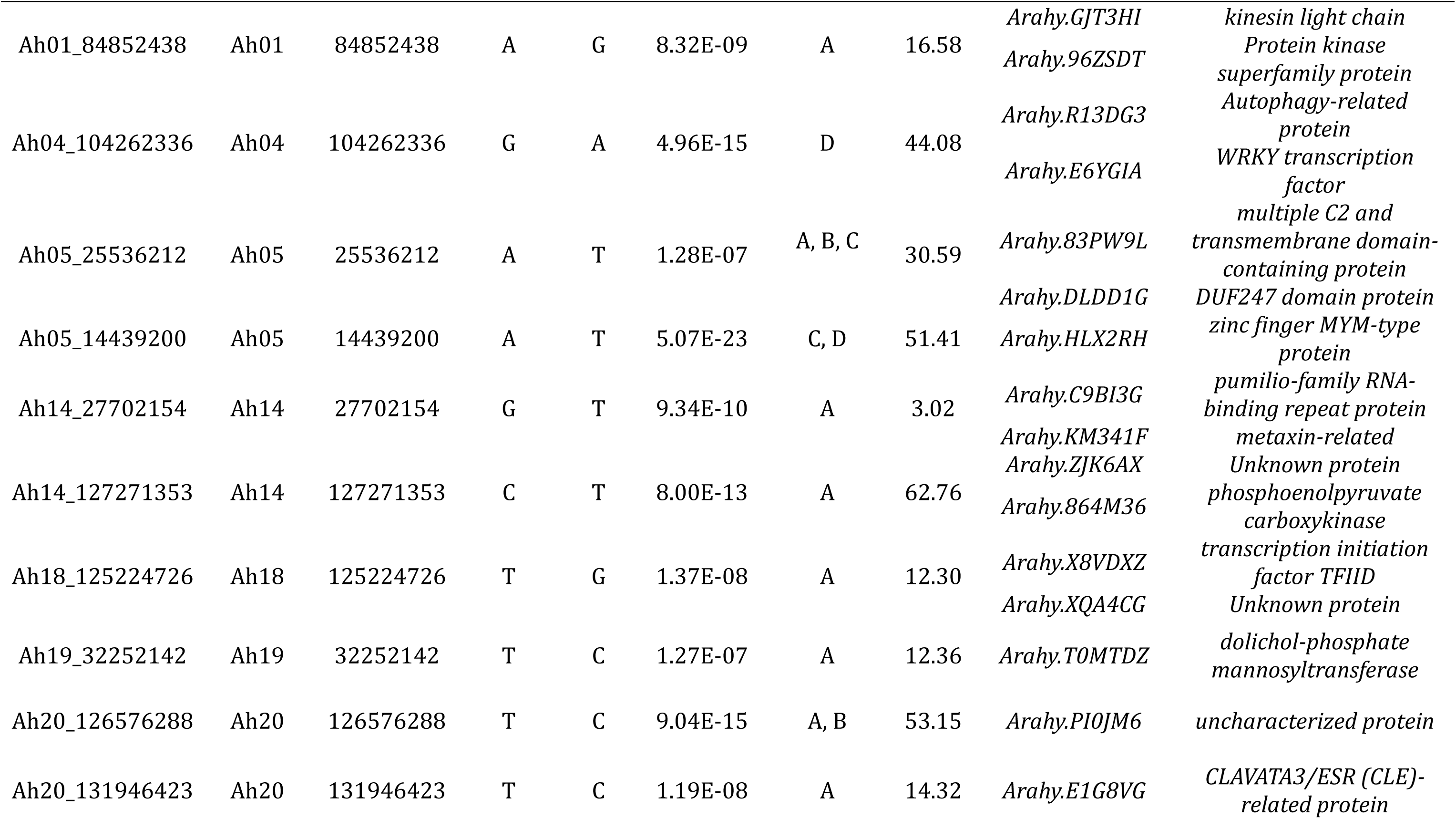
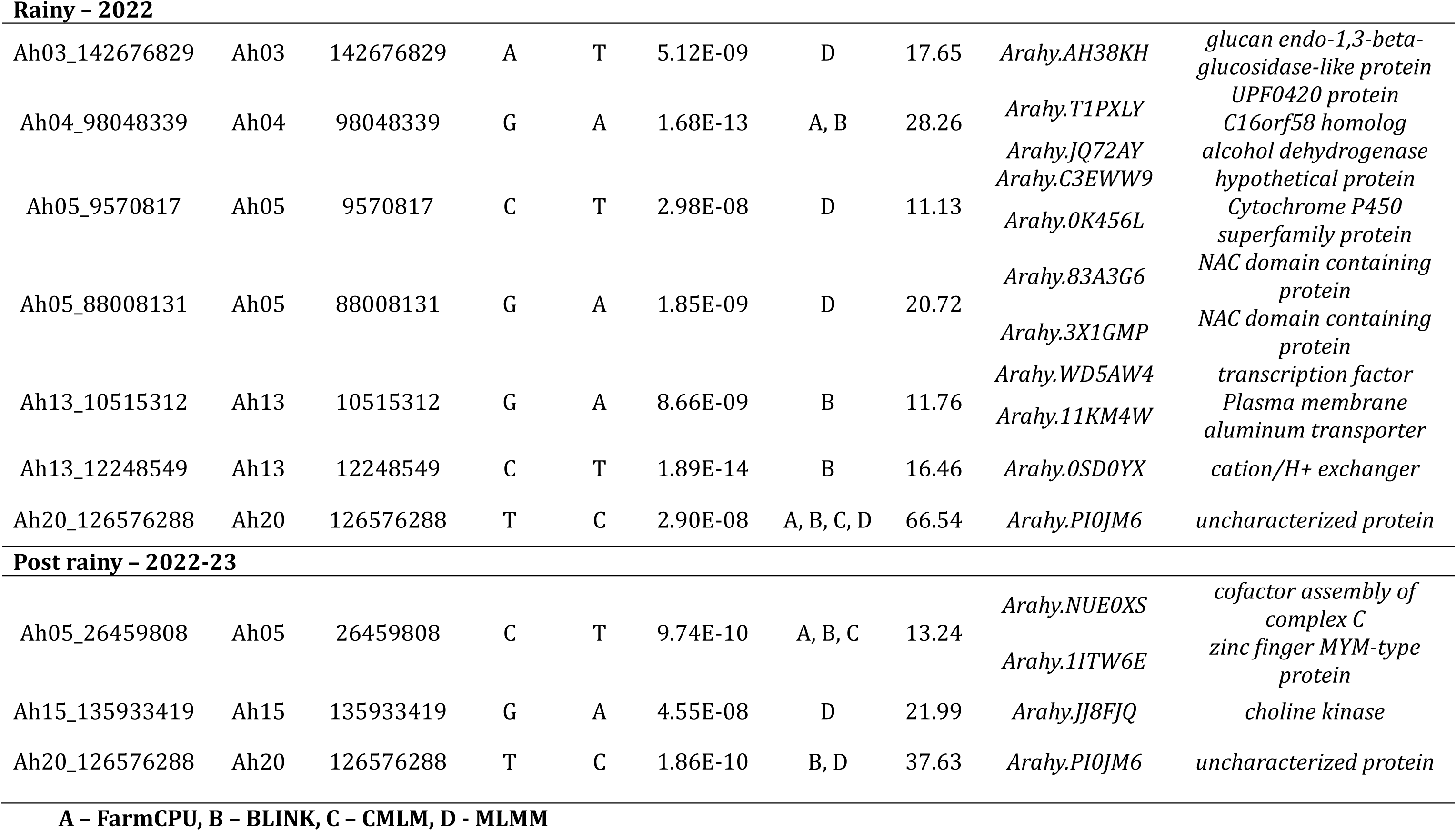
Marker trait associations and corresponding candidate genes identified for fresh seed dormancy.

**Figure 2.**
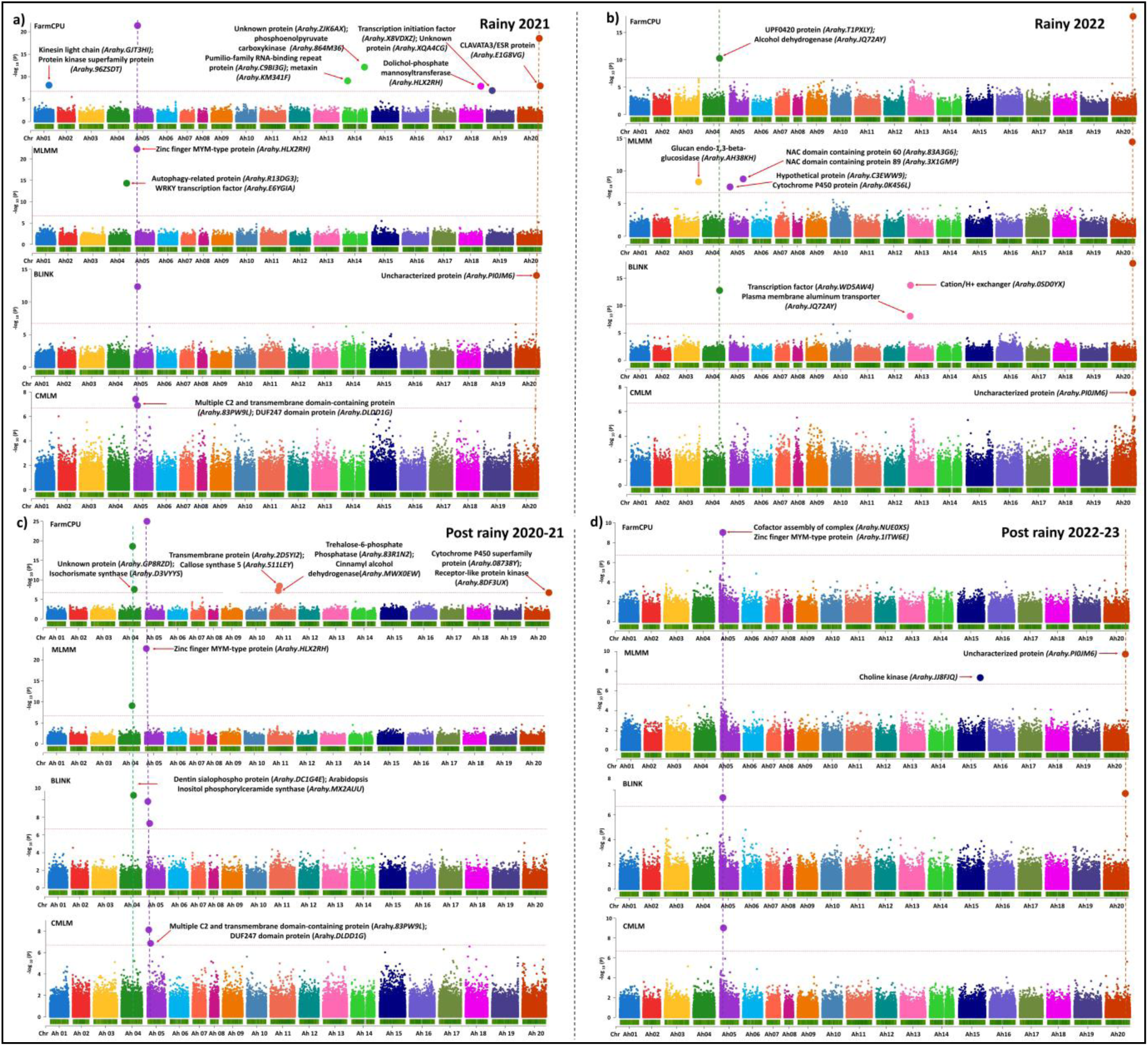
The Manhattan Plot represent the genome-wide association study (GWAS) results for groundnut days to 50% germination across four different models: FarmCPU, MLMM, BLINK, and CMLM. Data is shown for four growing seasons: Rainy 2021, Rainy 2022, and Post rainy 2020-21, Post rainy 2022-23. Each panel corresponds to a different model and illustrates the statistical significance of associations between genetic markers and days to 50% germination. **a)** Manhattan plot for Rainy 2021 **b)** Rainy 2022 **c)** Post rainy 2020-21 and **d)** Post rainy 2022-23.

Moreover, the pooled data from two rainy and post rainy seasons identified significant MTAs that were not detected in individual seasons. A total 12 MTAs were identified across six chromosomes, with PVE (%) from 6.04-71.94 (**Supplementary Table 2**). Interestingly, MTAs on Ah05 (Ah05_14439200) and Ah20 (Ah20_126576288), were consistently identified, in both pooled rainy and post rainy seasons. The MTA Ah20 (Ah20_126576288) was identified in both individual season analyses and pooled data (**Supplementary Figure 3a,b**). These consistent findings across multiple models and datasets strongly suggest the potential of these MTAs for further exploration. This led to an in-depth investigation of candidate genes underlying these MTAs.

### 3.3 Identification of candidate genes for fresh seed dormancy

Across four seasons, we identified 44 candidate genes associated with the trait, based on 27 distinct MTAs. Specifically, 7 MTAs were found in PR 2020–21, 10 in R 2021, 7 in R 2022, and 3 in PR 2022–23. These were involved in different cellular, molecular and biological functions related to the trait. These candidate genes encoded important protein like, *dentin sialophospho protein, Arabidopsis Inositol phosphorylceramide synthase, isochorismate synthase 2, zinc finger MYM-type protein, multiple C2 and transmembrane domain-containing protein, trehalose-6-phosphate phosphatase, callose synthase 5, receptor-like protein kinase 1, Protein kinase superfamily protein, Autophagy-related protein, probable WRKY transcription factor, Cytochrome P450 superfamily protein, alcohol dehydrogenase, NAC domain containing protein, transcription factor, Plasma membrane aluminium transporter, cation/H+ exchanger, cofactor assembly of complex C* and *choline kinase* were prominent genes known for their involvement in regulation dormancy/germination (**Table 1**).

Similarly, a total of 17 candidate genes were identified using 12 MTAs across both pooled seasons, specifically 9 genes in the pooled rainy season and 8 in the pooled post rainy season. The main proteins are *zinc finger MYM, proteasome subunit beta type-7-A, calcium-transporting ATPase, homeobox-leucine zipper protein, RING-box, 5-formyltetrahydrofolate cyclo-ligase, histone deacetylase, ninja-family protein* and *Target of Myb protein 1* (**Supplementary Table 2**). These both individual season and pooled seasons obtained genes are directly or indirectly involved in the regulation of dormancy/germination pathway.

In the 200kb genomic region, the identified Significant MTAs by individual seasons, total of 149 genes were extracted (Supplementary Table 3) of these, previously reported studies had identified 57 of the genes as candidate regulators of the process of seed dormancy or seed germination regulating *via* ABA, GA and ethylene signalling pathways (Supplementary Table 4). The genes corresponding to MTA on chromosome A05 (Ah05_9570817) in same season includes *Cytochrome P450 superfamily protein (Arahy.0K456L), glucan endo-1,3-beta-glucosidase (Arahy.B7TVVM),* also chromosome Ah20 in PR2020-21 having *Cytochrome P450 superfamily protein (Arahy.08738Y).* Similarly, *myb transcription factor (Arahy.2BK3KU; Arahy.S1DRQT; Arahy.1X4EZ2)* from Ah14_127271353, Ah20_131946423 and Ah13_12248549 are recognized as promising genes involved in ABA biosynthesis.

MTAs from pooled season results yielded a total of 57 genes (**Supplementary Table 5**). Previous studies have extensively reported 23 genes as potential genes among these included *serpin-ZX-protein (Arahy.GKKZ7Q), serine carboxypeptidase (Arahy.N7SB2Q)* on Ah05 and Ah08 from pooled rainy, similarly, from Ah04 having *F-box and associated interaction domain protein (Arahy.TJ6H4J)* were identified to regulate SD as reported in various crops (**Supplementary Table 6**). These genes were further analyzed to understand their expression patterns and functional roles in seed dormancy and germination.

### 3.4 Gene pathway for FSD regulated by ABA and GA

To explore the molecular mechanisms underlying dormancy and germination, we conducted an in-depth investigation into the functional roles of identified candidate genes involved in ABA and GA biosynthesis and signaling pathways. Among these candidates, the *Cytochrome P450 superfamily proteins (Arahy.08738Y*; *Arahy.0K456L)* were prominently recognized for their role in ABA catabolism. Additionally, several key players in ABA signaling were identified, including *WRKY family transcription factors (Arahy.E6YGIA)*, *protein kinase superfamily proteins (Arahy.96ZSDT)*, and *MYB transcription factors (Arahy.2BK3KU, Arahy.S1DRQT, Arahy.1X4EZ2)*. Furthermore, the *F-box protein interaction domain protein (Arahy.TJ6H4J)* and *GATA transcription factor (Arah.OIXS7Z)* emerged as a significant contributor to GA signaling. These findings highlight the intricate interplay between these hormones, providing valuable insights into the regulatory networks that govern seed dormancy and germination processes (**Figure 3**). To further validate the identified MTAs, we examined their polymorphism in a diverse panel of groundnut accessions.

**Figure 3.**
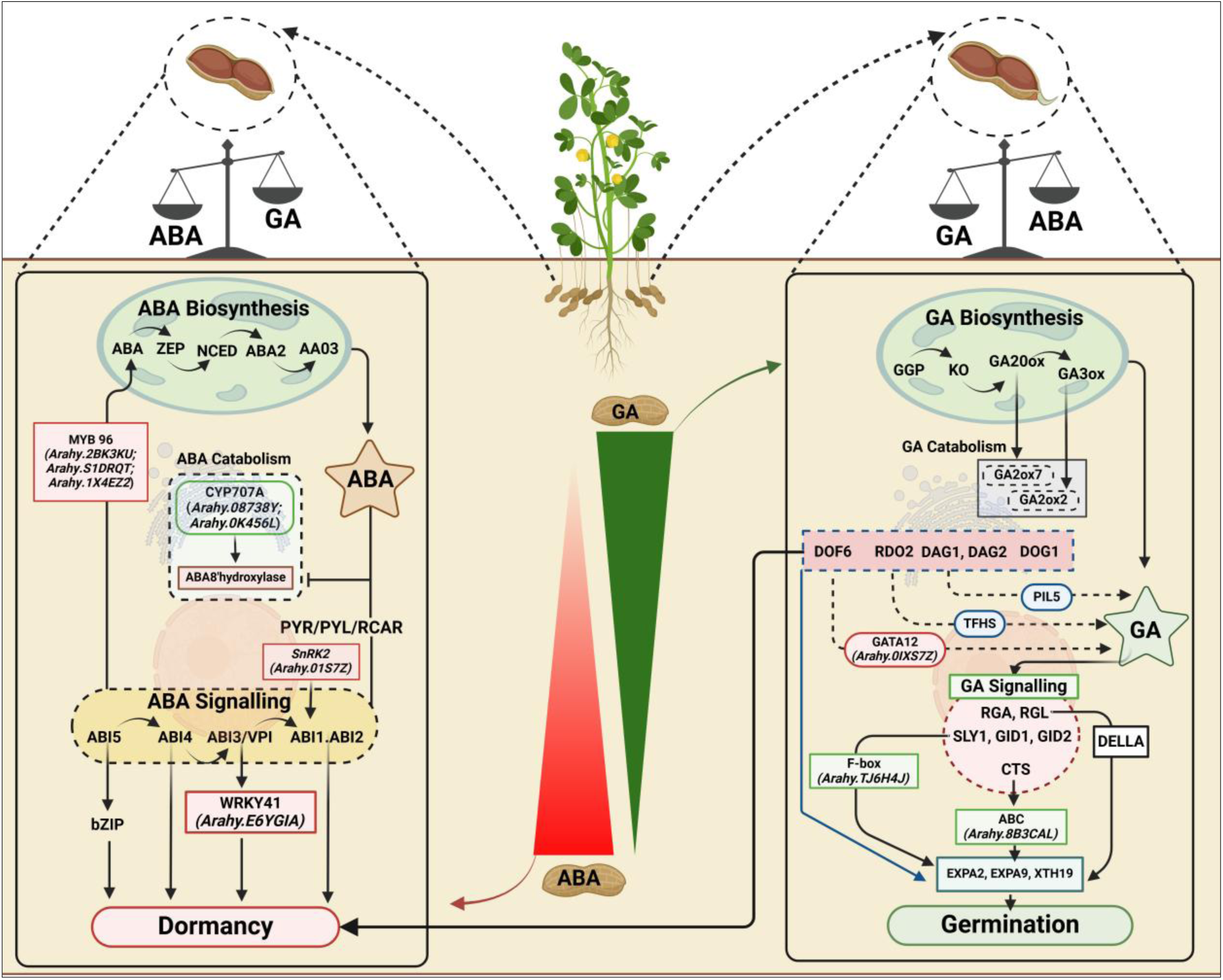
The diagram represents the interaction between the abscisic acid (ABA) and gibberellic acid (GA) pathways, which play crucial roles in regulating seed dormancy and germination. ABA biosynthesis and signaling pathways, highlighted on the left side, promote seed dormancy, with key genes like *ABI5*, *ABI4*, and *ABI3/VPI* involved in signaling, while genes like *MYB96* and *SnRK2* are central to the dormancy process. In contrast, GA biosynthesis and signaling pathways, shown on the right side, promote germination through genes like *RGA*, *RGL*, *SLY1*, and *GID1*, with direct pathway factors such as *DOF6* and *DOG1*. The balance between ABA and GA is depicted, illustrating how their opposing actions determine the seed’s transition from dormancy to germination.

### 3.5 Development and validation of KASP markers for FSD

In relation to the phenotyping data, a representative of the mini-core diversity panel, with varied durations of dormancy, was chosen to evaluate the robustness of the identified MTAs. The allele calls of the selected accessions, for 27 significantly associated SNP markers (identified season wise) and 12 associated SNPs (Pooled season), from WGRS and SNP array genotyping data based GWAS analysis were used for KASP assay design and validation. Among the 27 identified MTAs only 9 showed polymorphism between dormant and non-dormant accessions in the mini-core collection, specifically for markers Ah03_142676829, Ah04_79354879, Ah05_9570817, Ah05_88008131, Ah11_56153560, Ah13_12248549, Ah15_135933419, Ah20_141972519 and Ah20_126576288. Additionally, from a pooled analysis of 12 MTAs, 6 SNPs exhibited polymorphism Ah05_95779759, Ah08_17840995, Ah13_18020146, Ah15_68637668, Ah15_38301179 and Ah20_126576288 (**Supplementary Table 8**). Notably, the marker Ah20_126576288 consistently appeared across three growing seasons also in pooled seasons, indicating its reliability. So a total of 14 different markers showing clear polymorphism and 3 markers A09_29726644, B05_112344292 and B09_143731363 from the 58K panel were used for the validation.

KASP markers provide a cost-effective and efficient genotyping approach, enabling early-generation selection in breeding programs for indirect selection of target phenotypes [52]. To develop and validate diagnostic markers for FSD, 14 SNPs were targeted for KASP marker development, distributed across multiple chromosomes (three SNPs each on Ah05 and Ah15, two SNPs each on Ah13 and Ah20, and one SNP each on Ah03, Ah04, Ah08, and Ah11). Primers for these 14 SNPs were successfully designed and validated using a diverse validation panel categorized by dormancy period. The validation panel was comprised of non-dormant lines with dormancy period ranging from 1-7 days, moderate dormant lines ranging from 9-12 days and dormant lines ranging from 15-30 days of dormancy (**Supplementary Table 7**).

Among the 14 verified KASP markers, six markers (snpAH00577, snpAH00580, snpAH00582, snpAH00585, snpAH00586, snpAH00588) exhibited the expected polymorphism, clearly distinguishing between dormant and non-dormant genotypes (**Figure 4**). Additionally, three markers (snpAH00571, snpAH00572, and snpAH00573), located on chromosomes A09, B05, and B09, were included based on previous findings [37] (**Table 2**), further strengthening their relevance for FSD studies. In total, nine KASP markers demonstrated high polymorphism among non-dormant, moderate, and dormant groundnut genotypes. These markers serve as potential diagnostic tools for selecting dormant seeds and screening segregating breeding material at early stages of varietal development.

**Figure 4.**
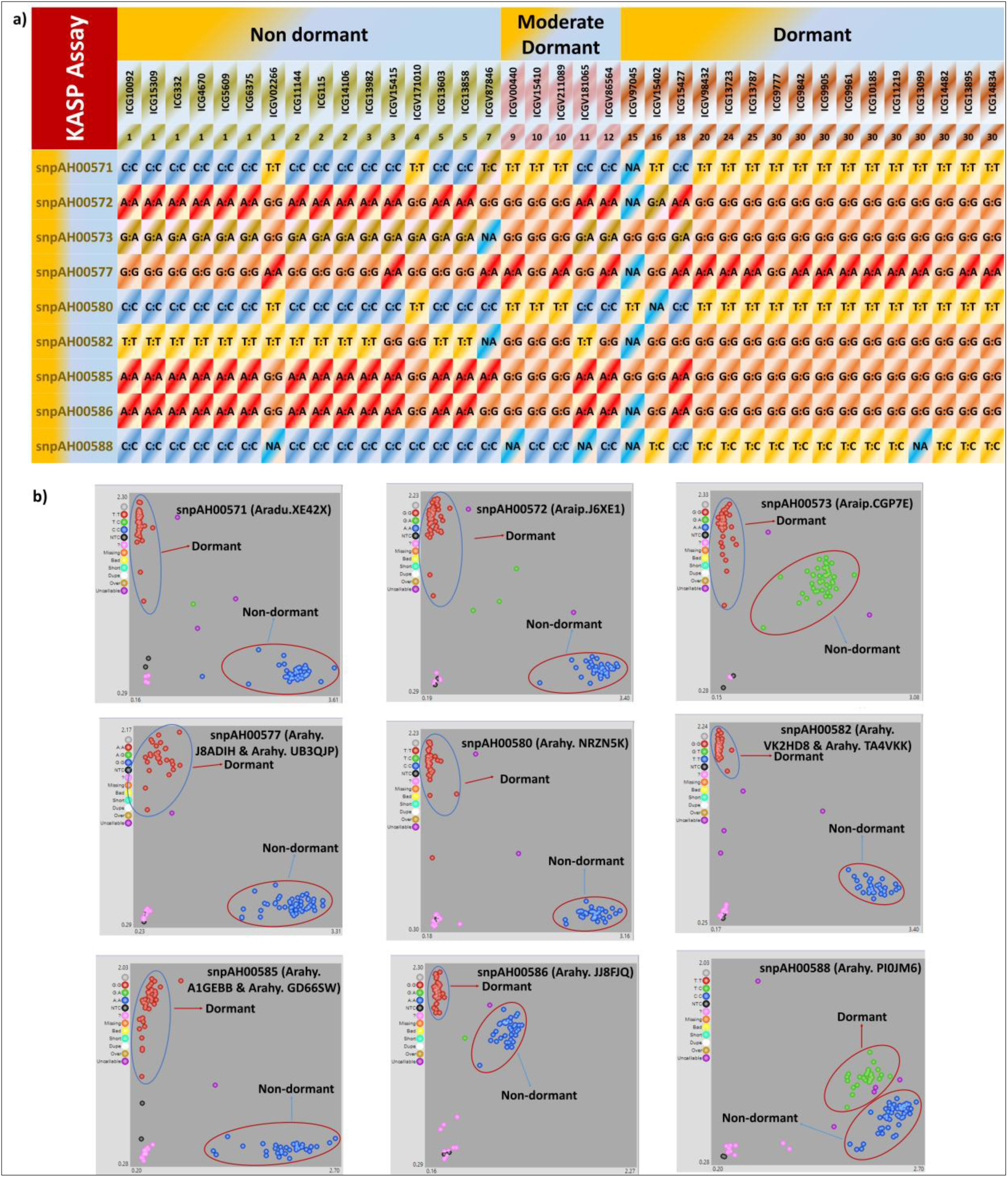
Development and validation of Kompetitive allele specific polymerase chain reaction (KASP) markers from potential candidate genes identified for fresh seed dormancy. (a) Validation panel includes 16 non-dormant lines (1-7days), 5 moderate-dormant (9-12 days) and 16 dormant lines (15-30 days) (b) snpAH00571, snpAH00572, snpAH00573 KASP markers from 58K “Axiom_*Arachis*” array and snpAH00577, snpAH00580, snpAH00582, snpAH00585, snpAH00586 and snpAH00588 KASP markers from WGRS were developed and validated for FSD.

**Table 2:** List of Validated KASP Markers from Candidate Genes Associated with Fresh Seed Dormancy in Groundnut.

| KASP ID | SNP ID | Chr.<br>No | Pos | Allele<br>(Ref) | Allele<br>(Alt) | Gene ID |
| --- | --- | --- | --- | --- | --- | --- |
| snpAH00571 | A09_29726644 | A09 | 29726644 | C | T | *Arahy.XE42X* |
| snpAH00572 | B05_112344292 | B05 | 112344292 | A | G | *Arahy.J6XE1* |
| snpAH00573 | B09_143731363 | B09 | 143731363 | A | G | *Arahy.CGP7E* |
| snpAH00577 | Ah05_95779759 | Ah05 | 95779759 | G | A | *Arahy.J8ADIH & Arahy.UB3QJP* |
| snpAH00580 | Ah08_17840995 | Ah08 | 17840995 | C | T | *Arahy.NRZN5K* |
| snpAH00582 | Ah13_18020146 | Ah13 | 18020146 | T | G | *Arahy.VK2HD8 & Arahy.TA4VKK* |
| snpAH00585 | Ah15_38301179 | Ah15 | 38301179 | A | G | *Arahy.A1GEBB & Arahy.GD66SW* |
| snpAH00586 | Ah15_135933419 | Ah15 | 135933419 | A | G | *Arahy.JJ8FJQ* |
| snpAH00588 | Ah20_126576288 | Ah20 | 126576288 | T | C | *Arahy.PI0JM6* |

### 3.6 Best allelic combination for achieving fresh seed dormancy

Among nine KASP markers validated from WGRS and the 58K ’Axiom_*Arachis*’ array (**Table 2**), only four markers; Ah05_95779759, Ah08_17840995, Ah15_135933419 and Ah20_126576288 exhibited the best allelic combination (A_1_T_1_G_3_Y), resulting in 24-30 days of dormancy. This combination includes all favourable alleles (A_1_T_1_G_3_Y) for dormancy, whereas the unfavourable allelic combination (G_4_C_1_A_3_C_2_) for these three markers led to only 1-6 days of dormancy, also When we combine the Ah05_95779759, Ah08_17840995, and Ah15_135933419 favorable allelic combination (A1T1G3), it gives 22-30 days of dormancy, while the unfavorable allelic combination (G4C1A3) gives 1-8 days of dormancy. Conversely, the other combinations for the remaining marker allelic combinations did not show significant variation in dormancy duration (**Figure 5**). The candidate genes associated with these three KASP markers were identified as *ATP-binding ABC transporter* (*Arahy.8B3CAL*) is involved in GA signalling, *calcium-transporting ATPase* (*Arahy.NRZN5K*), *uncharacterized protein (Arhay.PI0JM6)* involved in ABA pathway, and *choline kinase* (*Arahy.JJ8FJQ*) involved in both ABA-GA interaction.

**Figure 5.**
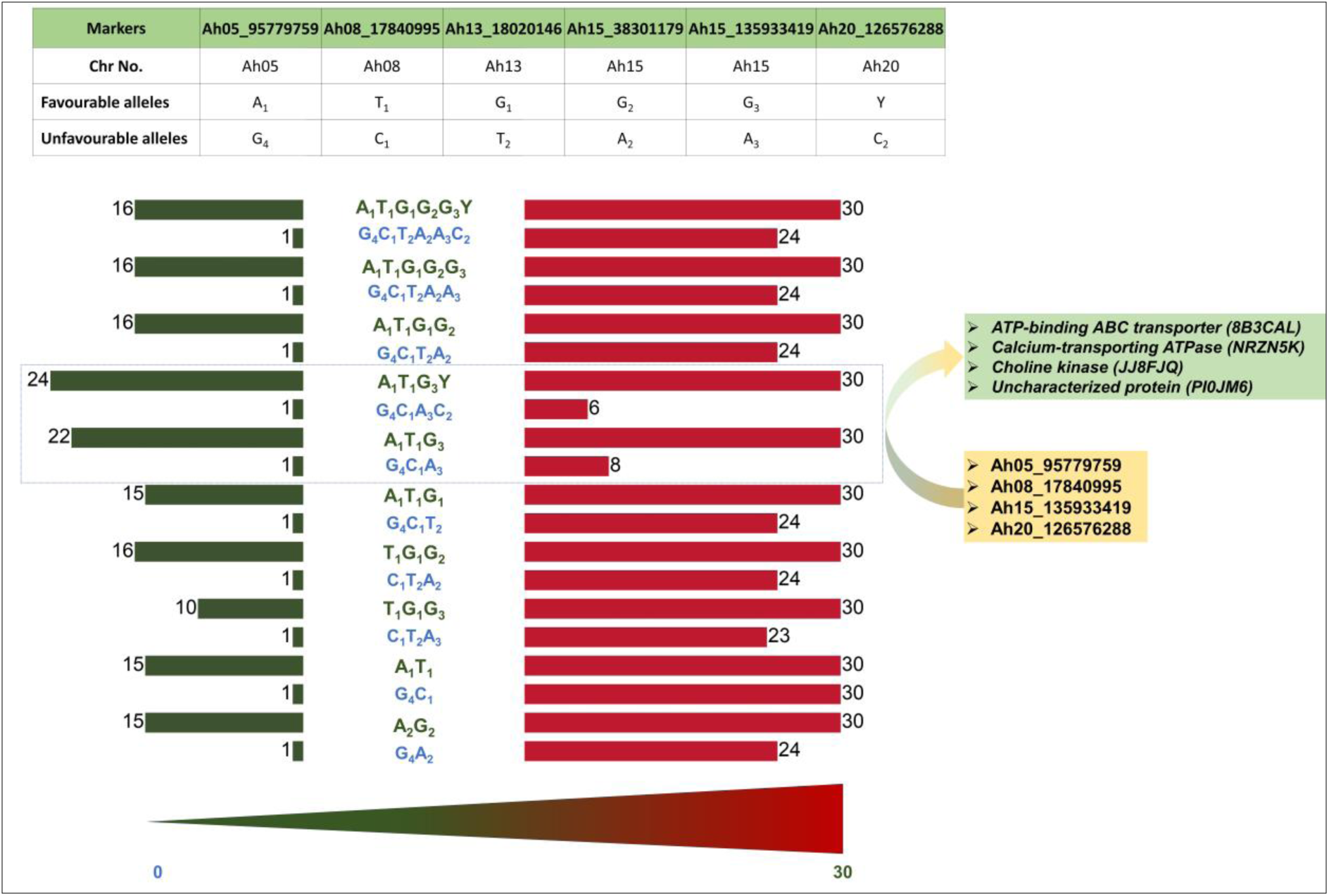
The diagram illustrates the relationship between specific allelic combinations of KASP markers and the duration of seed dormancy in days. Green letters represent favourable allelic combinations associated with longer dormancy periods (ranging from 24-30 and 22-30 days), with the “ A_1_T_1_G_3_Y” and “ A_1_T_1_G_3_” combination within the dashed blue rectangle on markers Ah05_95779759, Ah08_17840995, Ah15_135933419 and Ah20_126576288 showing the strongest association with extended dormancy (30 days). Conversely, blue letters indicate unfavourable allelic combinations, such as “G_4_C_1_A_3_C_2_” and “G_4_C_1_A_3_” for the same key markers, which result in significantly shorter dormancy (1-6 and 1-8 days). The associated candidate genes linked to these markers on chromosomes Ah05, Ah08, Ah15 and Ah20 are ATP-binding *ABC transporter, calcium-transporting ATPase*, *choline kinase*, and *uncharacterized protein* respectively, suggesting their potential roles in regulating the duration of dormancy.

## 4. Discussion

Groundnut is a widely cultivated leguminous crop valued for its high protein content and rich supply of unsaturated oils. However, one of the significant challenges in its production is PHS, where the seeds begin to germinate before harvest, leading to substantial yield losses. Genotypes exhibiting over 90% dormancy for a duration of 2 to 3 weeks are better adapted to regions with unpredictable rainfall patterns during the maturation stage. Incorporating such traits into breeding programs is crucial for ensuring stable yields under varying environmental conditions. GAB provides a significant advantage over traditional breeding methods by enabling the efficient tracking of desirable alleles in segregating populations through the use of trait-linked molecular markers [38]. In this study, a mini-core collection was analyzed using WGRS data and multi-season phenotyping to identify genomic regions and key candidate genes associated with FSD and germination traits. This integrative approach offers valuable insights into the genetic factors influencing FSD, which could contribute to the development of improved groundnut varieties with enhanced resistance to PHS.

GWAS has emerged as a robust method for identifying MTAs related to complex traits of interest. By leveraging genetic diversity within diverse germplasm/sets, GWAS has been widely utilized for uncovering genetic variation that influences various traits across crops[39]. Advanced GWAS models like CMLM, MLMM, FarmCPU, and BLINK in GWAS significantly improve the reliability and precision of identifying genetic associations in plant genomics [16]. CMLM is particularly effective at controlling for population structure and kinship, minimizing spurious associations [40]. On the other hand, MLMM addresses overfitting issues, making it particularly valuable for analysing complex traits. Both FarmCPU and BLINK are multi-locus models that boost statistical power by minimizing false positives and negatives. FarmCPU incorporate both fixed and random effects, making it highly effective across various trait analyses [40]. BLINK has computational efficiency, enabling rapid SNP detection while reducing false positives an essential feature when dealing with large genomic datasets [28,41,42]. These approaches can be further utilized to explore the genetic basis of complex traits, such as seed dormancy and germination, through resequencing-based genotyping and GWAS.

Resequencing-based genotyping reveals extensive natural variation, allowing for the exploration of functional genes and elite alleles within natural populations through GWAS. This approach enables the researcher to identify valuable genetic traits that can enhance breeding programs and improve crop performance [43,44]. By employing these models, we identified 27 significant MTAs for FSD trait through GWAS using a mini-core collection across individual seasons, and 10 significant MTAs in pooled seasonal results. Previous research findings reported two genomic regions on chromosomes B05 and A09 linked to FSD using QTL-Seq [45]. Also, a significant stable QTL linked with FSD was mapped on chromosome A04 [46]. Previously, we had deployed a 5K SNP assay in a bi-parental genetic mapping system and had identified five major QTLs on chromosomes Ah01, Ah06, Ah11, Ah16, and Ah17 in addition to two minor QTLs identified on Ah04 and Ah15 [47]. Notably, the *qFSD_A04-1* (119 Mb) was found near to *qPD_A04-2* [48], on chromosome A04, and *qFSD_B05-1* (112 Mb) was located near to the region of interest on chromosome B05 [45]. These findings highlight the reliability of GWAS in pinpointing significant MTAs in proximity to previously reported genomic regions.

Identification of candidate genes associated with QTLs/MTAs is provide insights into the molecular mechanisms regulating FSD. In particular, the roles of ABA, GA, and ethylene in regulating seed dormancy and germination have been well-documented across various crops. This suggests that ABA signalling, in conjunction with GA and ethylene interactions, plays a pivotal role in modulating both seed dormancy and the initiation of germination. In this study, genes identified through GWAS from both individual and pooled seasons were reviewed in existing literature to evaluate their functional roles in ABA and GA signalling pathways. This research elucidates how these genes contribute to the hormonal regulation of seed dormancy and germination processes, providing insights that could inform future breeding strategies to improve crop performance. *Cytochrome P450 (Arahy.08738Y)* in *Arabidopsis* encodes an enzyme known as *8’-hydroxylase*, which plays a crucial role in the catabolism of ABA. Studies have shown that mutants lacking the *cyp707a2* gene exhibit significantly elevated ABA levels up to six times higher than those found in wild-type plants leading to increased SD [49]. This suggests that *CYP707A2* functions as a negative regulator of SD by reducing ABA levels during imbibition. Additionally, transcript analysis of the *cytochrome P450 superfamily* genes revealed high expression levels in the non-dormant cultivar ICGV 91114, indicating their positive role in germination [47]. This evidence underscores the importance of *CYP707A2* and related *P450* enzymes in modulating ABA levels, thereby affecting seed dormancy and germination dynamics.

*MYB transcription factor (Arahy.2BK3KU; Arahy.S1DRQT; Arahy.1X4EZ2)* was identified in this study and this class of transcription factor is known for their role in ABA signaling [50]. *The MYB96 transcription factor* is thought to play a crucial role in regulating SD by promoting the biosynthesis of ABA through the enhancement of *NCED* genes, while simultaneously downregulating GA biosynthetic genes such as *GA20ox1* and *GA3ox1* in *Arabidopsis* [51]. In experiments, seeds from the *myb96-1* mutant germinated earlier than those with the wild-type *MYB96-1.* Conversely, seeds with an activated form of *myb96-1D* experienced delays in germination. This indicates that MYB96 is key in fine-tuning the timing of seed germination.

In this study, we identified a *protein kinase superfamily protein (Arahy.96ZSDT)* involved in ABA signalling. In *Arabidopsis*, two kinases that are specific to ABA signalling and related to seed dormancy and germination, *SNF1-RELATED PROTEIN KINASE 2.2 (SnRK2.2)* and *SnRK2.3*, are involved in ABA signalling [52]. Double mutants lacking both *snrk2.2* and *snrk2.3* show reduced expression of several genes that are typically induced by ABA, indicating their essential function in promoting ABA signalling. Additionally, redundant ABA-activated *SnRK2* kinases have been recognized as key regulators of seed maturation and dormancy in Arabidopsis [53]. This underscores the importance of *SnRK2 kinases* in the complex network of plant responses to environmental stresses and developmental cues. *SlWRKY37* is a *WRKY transcription factor* in tomato that plays a crucial role in regulating seed germination [48] revealed that the expression of *SlWRKY37* significantly decreases during the germination process.

Using CRISPR/Cas9 gene-editing technology, it was demonstrated that knocking out *SlWRKY37* enhances seed germination, whereas overexpressing this gene results in a delay of germination, playing an indirect role in dormancy. *WRKY family transcription factor family protein* (*Arahy.E6YGIA*) identified in this study was known to be involved in ABA signaling. In Arabidopsis, seeds that lack *WRKY41* show a marked decrease in the expression of *ABI3*, a gene vital for seed dormancy. Conversely, transgenic lines that overexpress *WRKY41* exhibit increased levels of *ABI3* expression [54]. Further analysis of the double mutant *wrky41 aba2* indicates that both *WRKY41* and ABA work together to regulate *ABI3* expression and seed dormancy. According to this synergy, *WRKY41* regulates *ABI3* crucially, impacting the dormant phase in seeds.

The *GATA-type zinc finger transcription factor family protein (Arahy.0IXS7Z)* identified in this study is involved in GA signalling. Furthermore, there is a novel interaction that involves *DOF6* and *RGL2*, which retains primary dormancy in *Arabidopsis* through *GATA transcription factors* [55]. The newly described *RGL2–DOF6* complex is required to activate *GATA12*, which is a gene that encodes for a *GATA-type zinc finger transcription factor* and is one of the downstream targets of *RGL2* expression in *Arabidopsis thaliana*. This offers a molecular basis for GA signalling repression, and correspondingly, that primary seed dormancy is maintained by expression [55–57].

The regulation of GA signalling gene *comatose* (CTS) is an important factor about dormancy/germination, and its activation induces germination through a peroxisomal protein of the *ATP-binding cassette (ABC) transporter* type [58,59]. The A*TP-binding ABC transporter (Arahy.8B3CAL)* and the *F-box interacting domain protein (Arahy.TJ6H4J)* represented two essential players in GA signalling. Regulation of the GA signalling gene *comatose* (CTS) is also an key component of whether seeds remain dormant or initiate germination. When activated, CTS facilitates the germination process by interacting with a peroxisomal protein from the *ATP-binding ABC transporter family* [58,59].

The SNP allele analysis of the diverse mini-core accessions for the identified significant MTAs indicate that accessions carrying all favourable dormant alleles tend to exhibit longer dormancy durations. The validation panel for FSD includes 40 genotypes representing 18 dormant, and 22 non-dormant genotypes. The deployment of diagnostic markers, including KASP markers, has become a routine practice in marker-assisted selection (MAS) for improving key agronomic traits in groundnut. These markers are widely used for resistance to nematodes, leaf rust, and late leaf spot, as well as for enhancing oil quality through high oleic acid content [38,60–66].

Recognizing the importance of FSD in groundnut breeding, we developed and validated six KASP markers (snpAH00577, snpAH00580, snpAH00582, snpAH00585, snpAH00586 and snpAh00588) located on chromosomes Ah05, Ah08, Ah13, Ah15 and Ah20 (**Table 2**). These markers effectively differentiate non-dormant, moderately dormant, and dormant groundnut genotypes, thereby facilitating MAS for dormancy. These markers target key candidate genes, enabling the efficient selection of dormant plants. Notably, one SNP from each associated region can be utilized for dormancy screening, improving the accuracy and effectiveness of breeding programs. Among these markers, snpAH00577, snpAH00580 snpAH00586 and snpAH00588 should be prioritized, as they are in the intergenic and downstream regions of *ATP-binding ABC transporter (Arahy.8B3CAL), calcium-transporting ATPase (Arahy.NRZN5K)*, *choline kinase (Arahy.JJ8FJQ)* and *uncharacterized protein (Arhay.PI0JM6)* respectively. These genes are known for their roles in regulating seed germination and dormancy pathways.

The *ATP-binding ABC transporter (Arahy.8B3CAL)* is directly involved in GA signalling (**Figure 3**), and *choline kinase* (*Arahy.JJ8FJQ*) plays a role in cell growth, development, and stress responses by maintaining membrane integrity under stress. Therefore, based on these findings, four KASP markers can be successfully used to identify dormant lines exhibiting 24-30 days of dormancy, surpassing the previously suggested in our studies Kumar et al. [45] and Bomireddy et al. [37], which could only detect lines with 2 weeks (14 days) of dormancy. These identified genes could be significantly regulating dormancy, enabling the lines to remain dormant for up to 30 days.

Our QTL-Seq study [45] also identified two genomic regions on chromosomes B05 and A09 associated with FSD and developed a potential marker (GMFSD1) on chromosome B05, which was successfully validated. Furthermore, we validated three markers (snpAH00571, snpAH00572, and snpAH00573), identified in our previous study [37] on chromosomes A09, B05, and B09, corresponding to GMFSD2, GMFSD3, and GMFSD4. Overall, chromosomes Ah05, Ah08, Ah15, Ah13 emerge as key FSD-associated regions along with Ah20. The validated KASP markers serve as robust diagnostic tools for selecting dormancy traits, providing valuable resources for marker-assisted breeding in groundnut. Ideally, 2-3 weeks of FSD period is sufficient. Therefore, accessions carrying a combination of dormant and non-dormant alleles for these markers can be effectively utilized to achieve the desired dormancy duration. These assays offer a targeted and efficient approach to selecting lines with optimized dormancy traits, significantly enhancing future molecular breeding initiatives in groundnut.

## 5. Conclusion

This study utilized WGRS data from a diverse mini-core collection, combined with phenotypic data collected over four seasons, to perform a GWAS analysis. The finding identified 27 MTAs in individual seasons and 12 MTAs pooled seasons for FSD. The analysis of these MTAs revealed potential candidate genes associated with FSD, highlighting the critical role of the ABA-GA balance in the indirect regulation of SD. Key proteins identified include *Cytochrome P450 superfamily proteins, protein kinase superfamily proteins, WRKY family transcription factors,* and *MYB transcription factors* involved in the ABA pathway, as well as *F-box interaction domain proteins, ATP-binding ABC transporters,* and *GATA-type zinc finger transcription factors* associated with the GA pathway were identified as key contributors to FSD regulation. Notably, our investigation into FSD revealed four highly effective KASP markers (snpAH00577, snpAH00580, snpAH00586 and snpAH00588) having allelic combination A_1_T_1_G_3_Y (Ah05_95779759, Ah08_17840995, Ah15_135933419 and Ah20_126576288) consistently resulted in a favorable 24-30 days of dormancy period, while the G_4_C_1_A_3_C_2_ combination significantly reduced dormancy to a mere 1-6 days, indicating its unfavorable impact. This study provides a foundation for the genetic improvement of groundnut seed dormancy, enabling the development of resilient cultivars to mitigate pre-harvest sprouting losses.

## Data availability statement

The phenotypic data used in this work provided Supplementary Table S1. The sequencing data generated in this study is deposited in NCBI with bio-project ID PRJNA1002116, PRJNA490835 and PRJNA490832.

## Institutional Review Board Statement

Not Applicable

## Informed Consent Statement

Not Applicable

## CRediT authorship Contribution Statement

**Manish K. Pandey:** Conceived the idea, supervised and finalized the manuscript. **Kuldeep Singh and Ramachandran Senthil:** Contributed seed material and in seed multiplication of the mini-core collection. **Pasupuleti Janila**: Provided the breeding lines for marker validation. **Deekshitha Bomireddy, Priya Shah, Vinay Sharma:** Phenotyped mini-core collection. **Sunil S. Gangurde, Namita Dube**: Performed the analysis. **D. Khaja Mohinuddin:** Performed the analysis and wrote the manuscript. **U. Nikhil Sagar**, **Hasan Khan, Tembhurne, V. Hanumanth Nayak, A. Amaregouda, Kisan Babu, Baozhu Guo**, **Boshou Liao**, **Rajeev K. Varshney:** Reviewed, edited and improved the manuscript. All authors have read and agreed to the published version of the manuscript.

## Acknowledgments

The authors are thankful to GeneBank, ICRISAT for their support in providing seed material and assistance in phenotyping work. D. Khaja Mohinuddin is grateful to ICRISAT for providing the facilities to conduct the work.

## Supplementary information

Table S1-Table S8

## Supplementary Figures

Figure S1-Figure S3

## References

[1] S.E.H. Taheri, M. Bazargan, P.R. Vosough, A. Sadeghian, A comprehensive insight into peanut: Chemical structure of compositions, oxidation process, and storage conditions, J. Food Compos. Anal 125 (2024) 105770.

[2] X. Wang, H. Zheng, Q. Tang, W. Mo, J. Ma, Effects of gibberellic acid application after anthesis on seed vigor of indica hybrid rice (*Oryza sativa* L.), Agronomy 9 (2019) 861.

[3] H.D. Upadhyaya, S.N. Nigam, Inheritance of Fresh Seed Dormancy in Peanut, Crop Sci. 39 (1999) 98–101.

[4] Y.B. Naganagoudar, P.V. Kenchanagoudar, S. Rathod, C.M. Keerthi, H.L. Nadaf, B.B. Channappagoudar, Inheritance of fresh seed dormancy in recombinant inbred lines (RIL) developed for mapping population TAG 24 x GPBD 4 in groundnut (*Arachis hypogea* L.), Legume Research-An International Journal 39 (2016) 844–846.

[5] M.K. Vishwakarma, M.K. Pandey, Y. Shasidhar, S.S. Manohar, P. Nagesh, P. Janila, R.K. Varshney, Identification of two major quantitative trait locus for fresh seed dormancy using the diversity arrays technology and diversity arrays technology-seq based genetic map in Spanish-type peanuts, Plant Breeding 135 (2016) 367–375.

[6] J.D. Bewley, Breaking down the walls—a role for endo-β-mannanase in release from seed dormancy?, Trends in Plant Science 2 (1997) 464–469.

[7] S.S. Parihar, M. Dadlani, S.K. Lal, V.A. Tonapi, P.C. Nautiyal, S. Basu, Effect of seed moisture content and storage temperature on seed longevity of hemp (*Cannabis sativa*), Indian J. Agri. Sci 84 (2014) 1303–1309.

[8] R. Finkelstein, W. Reeves, T. Ariizumi, C. Steber, Molecular Aspects of Seed Dormancy, Annu. Rev. Plant Biol. 59 (2008) 387–415.

[9] M.K. Pandey, A.K. Pandey, R. Kumar, C.V. Nwosu, B. Guo, G.C. Wright, R.S. Bhat, X. Chen, S.K. Bera, M. Yuan, H. Jiang, I. Faye, T. Radhakrishnan, X. Wang, X. Liang, B. Liao, X. Zhang, R.K. Varshney, W. Zhuang, Translational genomics for achieving higher genetic gains in groundnut, Theor Appl Genet 133 (2020) 1679–1702.

[10] R.K. Varshney, A. Bohra, J. Yu, A. Graner, Q. Zhang, M.E. Sorrells, Designing future crops: genomics-assisted breeding comes of age, Trends in Plant Science 26 (2021) 631–649.

[11] A. Gianinetti, P. Vernieri, On the role of abscisic acid in seed dormancy of red rice, Journal of Experimental Botany 58 (2007) 3449–3462.

[12] B. Mohanty, Genomic architecture of promoters and transcriptional regulation of candidate genes in rice involved in tolerance to anaerobic germination, Curr. Plant Biol. 29 (2022) 100236.

[13] E. Uffelmann, Q.Q. Huang, N.S. Munung, J. De Vries, Y. Okada, A.R. Martin, H.C. Martin, T. Lappalainen, D. Posthuma, Genome-wide association studies, Nature Reviews Methods Primers 1 (2021) 59.

[14] M.K. Pandey, H.D. Upadhyaya, A. Rathore, V. Vadez, M.S. Sheshshayee, M. Sriswathi, M. Govil, A. Kumar, M.V.C. Gowda, S. Sharma, F. Hamidou, V.A. Kumar, P. Khera, R.S. Bhat, A.W. Khan, S. Singh, H. Li, E. Monyo, H.L. Nadaf, G. Mukri, S.A. Jackson, B. Guo, X. Liang, R.K. Varshney, Genomewide association studies for 50 agronomic traits in peanut using the ‘reference set’comprising 300 genotypes from 48 countries of the semi-arid tropics of the world, PLoS One 9 (2014) e105228.

[15] M. Guo, L. Deng, J. Gu, J. Miao, J. Yin, Y. Li, Y. Fang, B. Huang, Z. Sun, F. Qi, W. Dong, Z. Lu, S. Li, J. Hu, X. Zhang, L. Ren, Genome-wide association study and development of molecular markers for yield and quality traits in peanut (Arachis hypogaea L.), BMC Plant Biol 24 (2024) 244.

[16] X. Liu, M. Huang, B. Fan, E.S. Buckler, Z. Zhang, Iterative usage of fixed and random effect models for powerful and efficient genome-wide association studies, PLoS Genetics 12 (2016) e1005767.

[17] C. Cui, H. Mei, Y. Liu, H. Zhang, Y. Zheng, Genetic diversity, population structure, and linkage disequilibrium of an association-mapping panel revealed by genome-wide SNP markers in sesame, Front. Plant Sci. 8 (2017) 1189.

[18] M.A. Rehman Arif, K. Neumann, M. Nagel, B. Kobiljski, U. Lohwasser, A. Börner, An association mapping analysis of dormancy and pre-harvest sprouting in wheat, Euphytica 188 (2012) 409–417.

[19] Q. Lu, X. Niu, M. Zhang, C. Wang, Q. Xu, Y. Feng, Y. Yang, S. Wang, X. Yuan, H. Yu, Y. Wang, X. Chen, X. Liang, X. Wei, Genome-wide association study of seed dormancy and the genomic consequences of improvement footprints in rice (*Oryza sativa* L.), Front. Plant Sci. 8 (2018) 2213.

[20] C. Li, P. Ni, M. Francki, A. Hunter, Y. Zhang, D. Schibeci, H. Li, A. Tarr, J. Wang, M. Cakir, J. Yu, M. Bellgard, R. Lance, R. Appels, Genes controlling seed dormancy and pre-harvest sprouting in a rice-wheat-barley comparison, Funct. Integr. Genom. 4 (2004) 84–93.

[21] S. Nakamura, F. Abe, H. Kawahigashi, K. Nakazono, A. Tagiri, T. Matsumoto, S. Utsugi, T. Ogawa, H. Handa, H. Ishida, M. Mori, K. Kawaura, Y. Orihara, H. Miura, A wheat homolog of MOTHER OF FT AND TFL1 acts in the regulation of germination, The Plant Cell 23 (2011) 3215–3229.

[22] A. Torada, M. Koike, T. Ogawa, Y. Takenouchi, K. Tadamura, J. Wu, T. Matsumoto, K. Kawaura, Y. Ogihara, A causal gene for seed dormancy on wheat chromosome 4A encodes a MAP kinase kinase, Curr Biol 26 (2016) 782–787.

[23] P. Kulwal, G. Ishikawa, D. Benscher, Z. Feng, L.-X. Yu, A. Jadhav, S. Mehetre, M.E. Sorrells, Association mapping for pre-harvest sprouting resistance in white winter wheat, Theor Appl Genet 125 (2012) 793–805.

[24] M. Lin, D. Zhang, S. Liu, G. Zhang, J. Yu, A.K. Fritz, G. Bai, Genome-wide association analysis on pre-harvest sprouting resistance and grain color in U.S. winter wheat, BMC Genomics 17 (2016) 794.

[25] A.S. Kaler, J.D. Gillman, T. Beissinger, L.C. Purcell, Comparing different statistical models and multiple testing corrections for association mapping in soybean and maize, Front. Plant Sci. 10 (2020) 1794.

[26] S.-B. Wang, J.-Y. Feng, W.-L. Ren, B. Huang, L. Zhou, Y.-J. Wen, J. Zhang, J.M. Dunwell, S. Xu, Y.-M. Zhang, Improving power and accuracy of genome-wide association studies via a multi-locus mixed linear model methodology, Sci Rep 6 (2016) 19444.

[27] B. Rakitsch, C. Lippert, O. Stegle, K. Borgwardt, A Lasso multi-marker mixed model for association mapping with population structure correction, Bioinformatics 29 (2013) 206–214.

[28] V. Segura, B.J. Vilhjálmsson, A. Platt, A. Korte, Ü. Seren, Q. Long, M. Nordborg, An efficient multi-locus mixed-model approach for genome-wide association studies in structured populations, Nat Genet 44 (2012) 825–830.

[29] Y.-M. Zhang, Z. Jia, J.M. Dunwell, The applications of new multi-locus GWAS methodologies in the genetic dissection of complex traits, Front. Plant Sci. 10 (2019) 100.

[30] H.D. Upadhyaya, P.J. Bramel, R. Ortiz, S. Singh, Developing a Mini Core of Peanut for Utilization of Genetic Resources, Crop Sci. 42 (2002) 2150–2156.

[31] M.K. Pandey, G. Agarwal, S.M. Kale, J. Clevenger, S.N. Nayak, M. Sriswathi, A. Chitikineni, C. Chavarro, X. Chen, H.D. Upadhyaya, M.K. Vishwakarma, S. Leal-Bertioli, X. Liang, D.J. Bertioli, B. Guo, S.A. Jackson, P. Ozias-Akins, & R.K. Varshney, Development and evaluation of a high density genotyping ‘*Axiom_ Arachis*’ array with 58 K SNPs for accelerating genetics and breeding in groundnut, Sci Rep 7 (2017) 40577.

[32] R. Li, C. Yu, Y. Li, T.-W. Lam, S.-M. Yiu, K. Kristiansen, J. Wang, SOAP2: an improved ultrafast tool for short read alignment, Bioinformatics 25 (2009) 1966–1967.

[33] P.J. Bradbury, Z. Zhang, D.E. Kroon, T.M. Casstevens, Y. Ramdoss, E.S. Buckler, TASSEL: software for association mapping of complex traits in diverse samples, Bioinformatics 23 (2007) 2633–2635.

[34] Y. Tang, X. Liu, J. Wang, M. Li, Q. Wang, F. Tian, Z. Su, Y. Pan, D. Liu, A.E. Lipka, E.S. Buckler, Z. Zhang, GAPIT Version 2: An Enhanced Integrated Tool for Genomic Association and Prediction, The Plant Genome 9 (2016) plant genome 2015.11.0120.

[35] A.M. Alqudah, J.K. Haile, D.Z. Alomari, C.J. Pozniak, B. Kobiljski, A. Börner, Genome-wide and SNP network analyses reveal genetic control of spikelet sterility and yield-related traits in wheat, Sci Rep 10 (2020) 2098.

[36] C. He, J. Holme, J. Anthony, SNP Genotyping: The KASP Assay, in: D. Fleury, R. Whitford (Eds.), Crop Breeding, Springer New York, New York, NY, 2014: pp. 75–86.

[37] D. Bomireddy, V. Sharma, S.S. Gangurde, D.K. Mohinuddin, R. Kumar, R. Senthil, K. Singh, M. Reddisekhar, S.K. Bera, M.K. Pandey, Multi-locus genome wide association study uncovers genetics of fresh seed dormancy in groundnut, BMC Plant Biol 24 (2024) 1258.

[38] R.K. Varshney, M.K. Pandey, P. Janila, S.N. Nigam, H. Sudini, M.V.C. Gowda, M. Sriswathi, T. Radhakrishnan, S.S. Manohar, P. Nagesh, Marker-assisted introgression of a QTL region to improve rust resistance in three elite and popular varieties of peanut (*Arachis hypogaea* L.), Theor Appl Genet 127 (2014) 1771–1781.

[39] R. Bohar, A. Chitkineni, R.K. Varshney, Genetic Molecular Markers to Accelerate Genetic Gains in Crops, Bio Techniques 69 (2020) 158–160.

[40] H. Tekeu, M. Jean, E.L. Ngonkeu, F. Belzile, Machine Learning-GWAS reveals the role of WSD1 gene for cuticular wax ester biosynthesis and key genomic regions controlling early maturity in bread wheat, bioRxiv (2023) 2023–11.

[41] X. Hu, J. Zuo, J. Wang, L. Liu, G. Sun, C. Li, X. Ren, D. Sun, Multi-locus genome-wide association studies for 14 main agronomic traits in barley, Front. Plant Sci. 9 (2018) 1683

[42] M. Adhikari, M.B. Kantar, R.J. Longman, C.N. Lee, M. Oshiro, K. Caires, Y. He, Genome-wide association study for carcass weight in pasture-finished beef cattle in Hawai’i, Front. genet 14 (2023) 1168150.

[43] Y. Zhao, H. Zhang, J. Xu, C. Jiang, Z. Yin, H. Xiong, J. Xie, X. Wang, X. Zhu, Y. Li, W. Zhao, M.A.R. Rashid, J. Li, W. Wang, B. Fu, G. Ye, Y. Guo, Z. Hu, Z. Li, Z. Li, Loci and natural alleles underlying robust roots and adaptive domestication of upland ecotype rice in aerobic conditions, PLoS Genetics 14 (2018) e1007521.

[44] J. Yu, W. Zao, Q. He, T.-S. Kim, Y.-J. Park, Genome-wide association study and gene set analysis for understanding candidate genes involved in salt tolerance at the rice seedling stage, Mol Genet Genomics 292 (2017) 1391–1403.

[45] R. Kumar, P. Janila, M.K. Vishwakarma, A.W. Khan, S.S. Manohar, S.S. Gangurde, M.T. Variath, Y. Shasidhar, M.K. Pandey, R.K. Varshney, Whole-genome resequencing-based QTL -seq identified candidate genes and molecular markers for fresh seed dormancy in groundnut, Plant Biotechnol. J 18 (2020) 992–1003.

[46] M. Zhang, Q. Zeng, H. Liu, F. Qi, Z. Sun, L. Miao, X. Li, C. Li, D. Liu, J. Guo, M. Zhang, J. Xu, L. Shi, M. Tian, W. Dong, B. Huang, X. Zhang, Identification of a stable major QTL for fresh-seed germination on chromosome Arahy. 04 in cultivated peanut (*Arachis hypogaea* L.), The Crop Journal 10 (2022) 1767–1773.

[47] D. Bomireddy, S.S. Gangurde, M.T. Variath, P. Janila, S.S. Manohar, V. Sharma, S. Parmar, D. Deshmukh, M. Reddisekhar, D.M. Reddy, P. Sudhakar, B.V.B. Reddy, R.K. Varshney, B. Guo, M.K. Pandey, Discovery of major quantitative trait loci and candidate genes for fresh seed dormancy in groundnut, Agronomy 12 (2022) 404.

[48] Z. Wang, J. Zhang, M. Gao, Q. Deng, Y. Zhang, M. Pei, Y. Zhao, Y.-D. Guo, H. Zhang, SlWRKY37 targets SlLEA2 and SlABI5-like7 to regulate seed germination vigor in tomato, Plant Physiol. Biochem 214 (2024) 108881.

[49] T. Kushiro, M. Okamoto, K. Nakabayashi, K. Yamagishi, S. Kitamura, T. Asami, N. Hirai, T. Koshiba, Y. Kamiya, E. Nambara, The Arabidopsis cytochrome P450 CYP707A encodes ABA 8′-hydroxylases: key enzymes in ABA catabolism, EMBO J 23 (2004) 1647–1656.

[50] F. Gosti, N. Beaudoin, C. Serizet, A.A. Webb, N. Vartanian, J. Giraudat, ABI1 protein phosphatase 2C is a negative regulator of abscisic acid signaling, The Plant Cell 11 (1999) 1897–1909.

[51] H.G. Lee, K. Lee, P.J. Seo, The Arabidopsis MYB96 transcription factor plays a role in seed dormancy, Plant Mol. Biol. 87 (2015) 371–381.

[52] H. Fujii, P.E. Verslues, J.-K. Zhu, Identification of two protein kinases required for abscisic acid regulation of seed germination, root growth, and gene expression in Arabidopsis, The Plant Cell 19 (2007) 485–494.

[53] K. Nakashima, Y. Fujita, N. Kanamori, T. Katagiri, T. Umezawa, S. Kidokoro, K. Maruyama, T. Yoshida, K. Ishiyama, M. Kobayashi, K. Shinozaki, K. Yamaguchi-Shinozaki, Three Arabidopsis SnRK2 protein kinases, SRK2D/SnRK2. 2, SRK2E/SnRK2. 6/OST1 and SRK2I/SnRK2. 3, involved in ABA signaling are essential for the control of seed development and dormancy, Plant and Cell Physiology 50 (2009) 1345–1363.

[54] Z.J. Ding, J.Y. Yan, G.X. Li, Z.C. Wu, S.Q. Zhang, S.J. Zheng, WRKY 41 controls Arabidopsis seed dormancy via direct regulation of *ABI 3* transcript levels not downstream of ABA, The Plant Journal 79 (2014) 810–823.

[55] P. Ravindran, V. Verma, P. Stamm, P.P. Kumar, A novel RGL2–DOF6 complex contributes to primary seed dormancy in Arabidopsis thaliana by regulating a GATA transcription factor, Molecular Plant 10 (2017) 1307–1320.

[56] A. Boccaccini, S. Santopolo, D. Capauto, R. Lorrai, E. Minutello, G. Serino, P. Costantino, P. Vittorioso, The DOF protein DAG1 and the DELLA protein GAI cooperate in negatively regulating the AtGA3ox1 gene, Molecular Plant 7 (2014) 1486– 1489.

[57] P. Stamm, P. Ravindran, B. Mohanty, E.L. Tan, H. Yu, P.P. Kumar, Insights into the molecular mechanism of RGL2-mediated inhibition of seed germination in Arabidopsis thaliana, BMC Plant Biol 12 (2012) 179.

[58] A. Skubacz, A. Daszkowska-Golec, Seed dormancy: the complex process regulated by abscisic acid, gibberellins, and other phytohormones that makes seed germination work, Phytohormones—Signaling Mechanisms and Crosstalk in Plant Development and Stress Responses (2017) 77–100.

[59] S. Footitt, Control of germination and lipid mobilization by COMATOSE, the Arabidopsis homologue of human ALDP, The EMBO Journal 21 (2002) 2912–2922.

[60] Y. Chu, C.L. Wu, C.C. Holbrook, B.L. Tillman, G. Person, P. Ozias-Akins, Marker-Assisted Selection to Pyramid Nematode Resistance and the High Oleic Trait in Peanut, The Plant Genome 4 (2011) 110–117.

[61] P. Janila, M.K. Pandey, Y. Shasidhar, M.T. Variath, M. Sriswathi, P. Khera, S.S. Manohar, P. Nagesh, M.K. Vishwakarma, G.P. Mishra, T. Radhakrishnan, N. Manivannan, K. Dobariya, R. Vasanthi, R.K. Varshney, Molecular breeding for introgression of fatty acid desaturase mutant alleles (ahFAD2A and ahFAD2B) enhances oil quality in high and low oil containing peanut genotypes, Plant Sci. 242 (2016) 203– 213.

[62] R.M. Kolekar, M. Sukruth, K. Shirasawa, H.L. Nadaf, B.N. Motagi, S. Lingaraju, P.V. Patil, R.S. Bhat, Marker-assisted backcrossing to develop foliar disease-resistant genotypes in TMV 2 variety of peanut (*Arachis hypogaea* L.), Plant Breeding 136 (2017) 948–953.

[63] S.K. Bera, J.H. Kamdar, S.V. Kasundra, P. Dash, A.K. Maurya, M.D. Jasani, A.B. Chandrashekar, N. Manivannan, R.P. Vasanthi, K.L. Dobariya, M.K. Pandey, P. Janila, T. Radhakrishnan, R.K. Varshney, Improving oil quality by altering levels of fatty acids through marker-assisted selection of ahfad2 alleles in peanut (*Arachis hypogaea* L.), Euphytica 214 (2018) 162.

[64] S.K. Bera, J.H. Kamdar, S.V. Kasundra, S.V. Patel, M.D. Jasani, A.K. Maurya, P. Dash, A.B. Chandrashekar, K. Rani, N. Manivannan, P. Janila, M.K. Pandey, R.P. Vasanthi, K.L. Dobariya, T. Radhakrishnan, R.K. Varshney, Steady expression of high oleic acid in peanut bred by marker-assisted backcrossing for fatty acid desaturase mutant alleles and its effect on seed germination along with other seedling traits, PLoS ONE 14 (2019) e0226252.

[65] D.B. Deshmukh, B. Marathi, H.K. Sudini, M.T. Variath, S. Chaudhari, S.S. Manohar, C.V.D. Rani, M.K. Pandey, J. Pasupuleti, Combining High Oleic Acid Trait and Resistance to Late Leaf Spot and Rust Diseases in Groundnut (*Arachis hypogaea* L.), Front. Genet. 11 (2020) 514.

[66] M.P. Jadhav, M.D. Patil, M. Hampannavar, Venkatesh, P. Dattatreya, K. Shirasawa, J. Pasupuleti, M.K. Pandey, R.K. Varshney, R.S. Bhat, Enhancing oleic acid content in two commercially released peanut varieties through marker-assisted backcross breeding, Crop Sci. 61 (2021) 2435–2443.

